# TroR is the primary regulator of the iron homeostasis transcription network in the halophilic archaeon *Haloferax volcanii*

**DOI:** 10.1101/2023.08.16.553580

**Authors:** Mar Martinez Pastor, Saaz Sakrikar, Sungmin Hwang, Rylee Hackley, Andrew Soborowski, Julie A. Maupin-Furlow, Amy Schmid

## Abstract

Maintaining intracellular iron concentration within the homeostatic range is vital to meet cellular metabolic needs and reduce oxidative stress. Previous research revealed that the haloarchaeon *Halobacterium salinarum* encodes four diphtheria toxin repressor (DtxR) family transcription factors (TFs) that together regulate the iron response through an interconnected transcriptional regulatory network (TRN). However, the metal specificity of DtxR TFs and the conservation of the TRN remained poorly understood. Here we identified and characterized the TRN of *Haloferax volcanii* for comparison. Genetic analysis demonstrated that *Hfx. volcanii* relies on three DtxR transcriptional regulators (Idr, SirR, and TroR), with TroR as the primary regulator of iron homeostasis. Bioinformatics and molecular approaches revealed that TroR binds a conserved *cis*-regulatory motif located ∼100 nt upstream of the start codon of iron-related target genes. Transcriptomics analysis demonstrated that, under conditions of iron sufficiency, TroR repressed iron uptake and induced iron storage mechanisms. TroR repressed the expression of one other DtxR TF, Idr. This reduced DtxR TRN complexity relative to that of *Hbt. salinarum* appeared correlated with natural variations in iron availability. Based on these data, we hypothesize that increasing TRN complexity appears selected for under variable environmental conditions such as iron availability.

## INTRODUCTION

Iron is essential for cellular processes and metabolic pathways, such as oxygen transport, respiration, the TCA cycle, gene regulation, and DNA synthesis (1). Consequently, iron can become life-limiting when unavailable. Conversely, excessive accumulation of intracellular iron results in oxidative stress damage, ultimately leading to cell death (2). Therefore, tight regulation of intracellular iron levels is critical for maintaining normal cellular growth and survival. Variations in iron availability can therefore drive niche expansion, adaptation, and ecological diversity of microbes (3,4).

Under stressful conditions such as iron starvation, the activation of transcriptional programs triggers specific metabolic pathways that enable the restoration of internal homeostasis, usually resulting in a delay in microbial growth (5,6). Such changes in the transcriptome occur through molecular systems called transcription regulatory networks (TRN) consisting of TFs (network nodes) that bind to specific cis-regulatory sequences (edges) to differentially regulate sets of target genes (downstream nodes). TFs work hierarchically to determine what mRNA transcripts must be synthesized, when, and at what level by binding specific cis-regulatory sequences within promoter regions (7,8). The transcriptional response governed by TRNs thereby facilitates dynamism in response to environmental stimuli, and the complexity of the interaction between components determines the context-specific pattern of the gene expression (9–11). Archaea provide an excellent model system for studying TRNs because of their relatively simple genome organization. Archaeal transcriptional machinery core components share relationship to eukaryotes, including TATA-binding protein (TBP) and transcription factor B (TBP); however, the modulation of gene expression by TFs is closely related to those in bacteria (10,12). Furthermore, archaeal genes are relatively small (averaging less than 1 kb in length) and are organized in operons transcribed as polycistronic mRNAs. Operons are separated by short (less than 200 bp) non-coding regulatory regions, resulting in high gene density (13).

Bacteria and archaea have evolved a variety of mechanisms to regulate intracellular iron concentrations. In iron poor conditions, uptake systems encoded by the *iucABC-bdb-dat* gene cluster synthesize and transport siderophores (14). These secreted low molecular weight compounds bind ferric iron with high affinity, then are actively imported to deliver iron intracellularly. Iron storage mechanisms encoded by the highly conserved gene *dpsA* are employed to protect the cell from free radical damage (15,16). In low iron conditions, iron-sulfur clusters, important co-factors in proteins essential in cellular processes such as respiration, can be damaged and require re-synthesis by the SUF system (17). Iron homeostasis mechanisms are regulated by transcription factors (TFs) that cluster to either the ferric uptake regulatory (Fur, PF01475) or diphtheria toxin repressor (DtxR, PF0135) superfamilies. Fur TFs regulate iron homeostasis in Gram-negative bacteria and Gram-positive bacteria with low GC content (5), while the transition metal ion activated DtxR TFs are identified to be central regulators of iron metabolism in Gram-positive bacteria (18). DtxR is widely conserved across bacteria and archaea (4,9,15,19,20), while Fur appears more commonly in bacteria (PF01475 showed 27,092 hits in bacterial genomes versus 275 in archaeal genomes). In particular, Leyn and Rodionov showed that the DtxR family of TFs is differentially expanded across Archaea (19). Among the 26 archaeal species analyzed belonging to 3 different phyla (Euryarchaeota, Crenarchaeota, and Korarchaeota) and adapted to variable natural environments, two to six DtxR paralogs were identified per genome with an average of 3.3 DtxR members per species.

Many archaea are found in harsh environments, enabling direct tests of the hypothesis that highly interconnected TRNs facilitate rapid adaptation to fluctuating environmental conditions (6,10,21). *Halobacterium salinarum*, an organism classified to the Euryarchaeota phylum, class Haloarchaea serves as a prime illustration of an organism capable of surviving to significant iron fluctuations. Despite its reliance on iron for growth, this haloarchaeon exhibits remarkable resilience to conditions of severe iron deficiency, enduring periods lasting for months without iron availability (22). Specifically, *Hbt. salinarum* uses TRNs to adapt to rapidly changing iron environments such as the water column of closed, evaporative saturated salt lakes and ponds that fluctuate with environmental conditions (e.g. the Great Salt Lake, Utah)(23–25). Our prior research demonstrated that *Hbt. salinarum* expresses four interrelated DtxR paralogs (Idr1, Idr2, SirR, and TroR) that are directly involved in the response to iron stress, as mutations in each paralog showed impairments in growth and intracellular iron accumulation (9,26). The TRN established between these four DtxR TFs is an interlocking loop of transcriptional regulation that facilitates dynamic changes to tune quick responses in response to changes in iron availability (9). It is still unknown whether this complex TRN structure is conserved with other species of the Haloarchaea.

To address this question, *Haloferax volcanii*, a hypersaline archaeon related to *Hbt. salinarum,* was used here as a model organism to study and compare the transcriptional mechanism of adaptative response to iron imbalance*. Hfx. volcanii* is an extreme halophile first isolated from Dead Sea sediment, a relatively stable environment where ferric iron is bioavailable (27–29).

*Hfx. volcanii* has been used in the laboratory as a model organism since the late 1980s when tools for genetic and molecular manipulation were developed (30,31). *Hfx. volcanii* grows optimally in lower sodium and higher magnesium concentrations than *Hbt. salinaum* in the laboratory and natural environment. Both use a salt-in strategy to counterbalance the high osmolarity of the medium. In both organisms, iron detoxification systems are conserved with bacteria, including siderophore biosynthesis, rudimentary SUF systems encoded by *sufBCD* genes, and DpsA-dependent iron storage (17,20,32). These systems are regulated by the DtxR TRN in *Hbt. salinarum* (9,26). Here we inquire if the structure and function of the DtxR TRN of *Hfx. volcanii* is conserved with that of *Hbt. salinarum* given their different hypersaline ecological niches.

Using a combination of genetic analysis, functional genomics, and promoter-reporter assays, here we develop a model for DtxR TRN structure and function in regulating iron homeostasis in *Hfx. volcanii*. Our model demonstrates that *Hfx. volcanii* expresses three homologous DtxR proteins (TroR, Idr, and SirR) but mainly utilizes TroR, an essential TF, to regulate the expression of the iron regulon. This points towards the hypothesis that in steady iron conditions, only a simplified TRN is required to control intracellular iron ions within a healthy range. The data in this study together suggest that, even with the extensive evolutionary expansion of the DtxR TF family encoded in genomes across the halophile phylogeny, the selective pressure of the different habitats shapes different interactions between those TFs and their target genes, enabling adaptative physiology to improve fitness.

## MATERIALS AND METHODS

### Strains and plasmids

All *Hfx. volcanii* strains originated from the wild-type strain DS2 (30) (maintained in the Maupin-Furlow and Schmid labs). The parent strain (H26) resulted from the deletion of the gene *pyrE2*, resulting in auxotrophy for uracil (Δ*pyrE*), allowing for selection of deletion mutants. Deletion mutants Δ*idr* (HVO_0538), Δ*sirR* (HVO_0819), Δ*troR* (HVO_0863), and Δ*arsR* (HVO_1766) were constructed in the H26 background by “pop-in / pop-out” homologous recombination (30). Briefly, flanking regions of ∼500 nt upstream and downstream of the gene of interest were amplified by PCR using specific primers designed to include HindIII and XbaI restriction sites (31) and parent H26 DNA as a template. The PCR product was cloned into HindIII-XbaI linearized plasmid pTA131, resulting in pJAM3194, pJAM3198, pJAM3196 and pAKS188 for deletion of *troR*, *sirR, idr*, and *arsR*, respectively. These plasmids were transformed into H26 as described in the Halohandbook (https://haloarchaea.com/halohandbook/) resulting on the strains HV173, HV174, HV175, and HV428. All primers, plasmids, and resultant strains are listed **Table S1**. Homologous recombination and two successive rounds of selection on novobiocin (1 ng·ml^-1^) and 5-fluoroorotic acid (50 µg·ml^-1^) were used to delete the genes of interest. HA-tagged strains were constructed by inserting the hemagglutinin epitope tag (-HA) in frame with the 3’ end of the gene of interest generating the strains HV284 (*troR-HA*), HV176 (*sirR-HA*), HV177 (*idr-HA*), and HV432 (*arsR-HA*). PCR, Sanger sequencing, whole genome resequencing (WGS), and western blot were used to verify the cloning, the mutations, and the absence of the translated protein. For *Escherichia coli* recombinant DNA experiments, DH5α competent cells were prepared and transformed as described previously (33).

### Media recipes and growth conditions

For *E. coli* cultures used for cloning, cells were grown overnight in Luria-Bertani medium broth (LB) supplemented with carbenicillin (50 µg.ml^-1^) for plasmid acquisition selection (34). In all cases, 2 % (w/v) agar from Difco agar was added to media recipe for solid media plate preparation. To routinely grow *Hfx. volcanii,* cells were refreshed from −80°C stock on rich media YPC18% plates [2.5 M NaCl, 50 mM KCl, 100 mM MgSO_4_·7H_2_O, 100 mM MgCl_2_·6H_2_O, 40 mM TrisHCl pH 7.5, 6.5 % (w/v) yeast extract from Fisher Scientific, 0.1 % (w/v) peptone from Oxoid, 0.1 % (w/v) casamino acids from VWR, 2 % (w/v) agar]. The following reagents were added to YPC18% after autoclaving once the preparation had cooled: 3 mM CaCl_2_ and uracil 50 µg.ml^-1^. Plates were incubated at 42°C until colonies were visible (48-72 h). For each experiment, three single colonies of each strain were pre-inoculated in 3 ml liquid-rich media YPC18% until saturation in a shaking incubator at 225 rpm and 42°C overnight.

To compare cell growth under varying concentrations of iron, a minimal media formulation (PR) was modified from previous study (35). The modified PR contained: 3 M NaCl, 100 mM KCl, 290 mM MgSO_4_·7H_2_O, 12 mM Tris HCl pH7.5, 5 mM NH_4_Cl, 0.5 mM K_2_HPO_4_, 0.8 ml thiamine at 1 mg·ml^-1^, 0.1 ml biotin at 1 mg·ml^-1^, 5 mM NaHCO_3_, 0.5% (w/v) glucose. All salts added to the formulated minimal media were metal-free (NaCl Beantown Chemical, 99.999% purity; KCl Alfa Aesar Puratronic, 99.999% purity; MgSO_4_·7H_2_O Alfa Aesar Puratronic, 99.997% purity; NH_4_Cl Beantown Chemical, 99.999% purity). The medium was treated with 10 g·l^-1^ of BT Chelex 100 resin (BioRad, Hercules, CA) for 2 h and then filtered using a vacuum-driven 0.22 µm filter (Olympus plastics, Genesee Scientific) prior to the inoculation of cells to remove any remaining trace of metal. PR was rendered metal replete (PR+) by adding the following metals in concentrations of 1.5 µM ZnSO_4_, 1.8 µM MnSO_4_, 0.1 µM CuSO_4_, 100 µM FeSO_4_ (all metals from Alfa Aesar, >99.99% purity). For PR-, the same media formulation was also treated with 100 µM BPS (bathophenanthroline disulfonate) iron chelator overnight and vacuum filtered prior to adding Zn, Mn, and Cu at the same concentrations as PR+, but with iron omitted. For all experiments comparing cells grown with and without iron, YPC pre-inoculation cultures (described above) were pelleted at 6,000 x g, resuspended in PR+, and grown overnight. The next morning, cells were washed once with PR-and inoculated to an initial OD600 ∼ 0.1 in PR+ or PR-. Resuspension volumes varied by experiment: 200 µl for growth curve measurements in the Bioscreen plate reader (described below), 3 ml culture for promoter-reporter experiments (see information below), or 50 ml for RNA extraction (RNA-seq or RT-qPCR) and ChIP experiments (ChIP-seq and ChIP-qPCR).

### Phylogenetic analysis

To detect DtxR homologs across archaea and build the DtxR phylogenetic tree, we used the web application AnnoTree (http://annotree.uwaterloo.ca/app/) with KEGG identifier K03709 (DtxR family transcriptional regulator) and all available archaeal genome sequences as input, accessed August 14, 2023 (36). AnnoTree default parameters were used as criteria for DtxR ortholog inclusion in the tree: minimum 30% sequence identity, E-value maximum of 1 x 10^-5^, minimum 70% subject and query alignment. DtxR sequence identity amongst TroR, Idr, and SirR was detected by reciprocal protein BLAST with default parameters. Multiple Sequence Alignments between TroR, SirR, and Idr of *Hfx. volcanii* was conducted using Clustal Omega with default parameters.

### Growth rate assays and analysis

To quantify cell growth rates in response to varying concentrations of iron, *Hfx. volcanii* cells were inoculated in the previously described media (PR+ and PR-) in 2×100 well-multiplates and incubated at 42°C with continuous shaking at 225 rpm in a Bioscreen C microbial growth analyzer (Growthcurves USA, Piscataway, NJ, USA) with OD measurements taken every 30 min. Growth of each strain was measured in 2-18 replicates as specified in the figure legend. For the titration experiment (Figure 1 A and B), variations in the iron concentrations are specified in the figure and text. Raw growth data were fitted using logistic regression, and growth metrics derived from fitted curves were calculated using the growthcurver package in the R coding environment (37). Growth metrics include the area under the logistic fitted curve (AUC) and the maximum instantaneous growth rate (µ).

**Figure 1.**
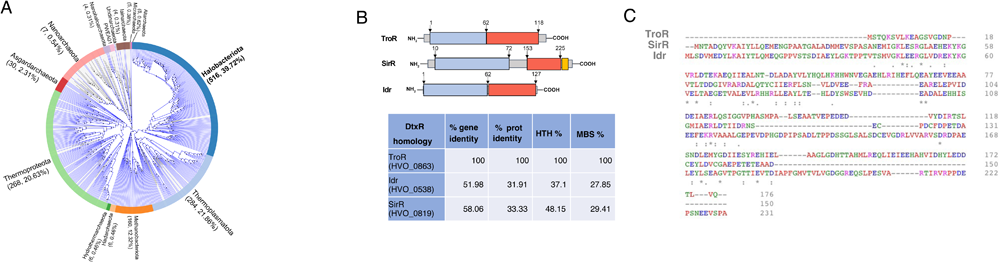
DtxR family of proteins is conserved across archaea. (A) The phylogenetic tree of the DtxR family of proteins has been performed using the AnnoTree tool for visualization of the annotated microbial tree of life (36). Each colored band and corresponding annotation show phylum-level assignments, and tree resolution represents the genus level. Grey lines indicate the genera included in the analysis, and blue highlighting shows the genera encoding DtxR orthologs. Parenthetical numbers next to each phylum name reports the number of genomes detected in that phylum, then the percentage of representative genomes that encode DtxR orthologs. (B) Diagram at top represents the DtxR protein structure for *Hfx. volcanii* paralogs. Numbers above the protein structure diagrams correspond to amino acid residue numbers. The conserved N-terminal region helix-turn-helix (HTH) domain is involved in DNA binding (blue region) and the C-terminal domain in metal binding (red region). One-third of the DtxR family proteins also contain the conserved domain SH3 in the C-terminus (yellow), predicted to control protein-protein interaction. Table at bottom reports the amino acid sequence identity of each DtxR paralog to that of TroR. (C) ClustalOmega alignment of the three proteins from *Hfx. volcanii* shows the specific conserved amino acids marked with asterisks. Dots mark residues that are conserved in two of the three proteins and dashes represent gaps in the alignment.

### Whole genome re-sequencing and analysis of deletion strains

Cells (2 mL) grown in YPC rich media (OD ∼ 0.9) were pelleted by centrifugation at 16,000 x *g* for 1 min at room temperature (rt). The supernatant was removed by pipetting out the residual liquid. Cells were lysed by adding 200 μl of DNase-free water and resuspending them up and down using a 3 ml syringe and 0.5 mm needle (BD Biosciences). An equal volume of phenol:chloroform:isoamyl alcohol (25:24:1) was added to the cell suspension to separate the genomic DNA (gDNA). Organic and aqueous phases were mixed by thorough inversion of the tubes and then separated by centrifugation (5 min 16,000 x *g*, rt). The upper aqueous phase that contains the nucleic acid suspension was transferred to a new 1.7 mL Eppendorf tube and 1/10 (v/v) of 3 M sodium acetate pH 5.2 was added. Finally, the gDNA was precipitated by adding 2X volume of 100% EtOH and incubated overnight at −20°C. The precipitated gDNA was pelleted by centrifugation (21,000 x *g*, 10 min at 4^°^C) in a tabletop centrifuge (Model 5424, Eppendorf). The supernatant was discarded by decantation. The pelleted gDNA was washed by adding 250 μL ice-cold 70% EtOH and centrifuged (21,000 x *g*, 5 min at 4^°^C). The residual liquid was discarded by decantation, and the gDNA pellet was air-dried for 5 min. gDNA was finally resuspended on 200 μL of DNase-free water. The obtained gDNA was sonicated using the Bioruptor Sonication System (Diagenode) to mechanically break the DNA by ultrasound. 15 cycles of 1 min each (30 sec on / 30 sec off) were enough to result in an optimal distribution of DNA fragment sizes for sequencing (300-800 bp). Samples were sequenced at the Duke Sequencing and Genomic Technologies core facility. There, library preparation was conducted using the KAPA HyperPrep Library Kit (Roche) and the whole genome was sequenced by Illumina HiSeq4000. The reads were mapped to the *Hfx. volcanii* DS2 genome (GenBank assembly GCA_000025685.1) and analyzed for mutations using the *breseq* package with the default parameters (38), as previously described . Raw sequencing data are available via the NCBI Sequence Read Archive (SRA) accession PRJNA1003915. Breseq mutation calls are listed in **Table S1**.

### Quantitative reverse transcriptase PCR (RT-qPCR)

Total RNA was extracted from 50 mL cultures grown in PR+ or PR-as described above. Aliquots of each culture (10 mL, OD_660_ of 0.3-0.5 mid-log) were harvested by centrifugation (5,000 x *g*, 5 min at 4°C) (Eppendorf centrifuge 5430R). The supernatant was immediately discarded. The pellets were snap-frozen in liquid nitrogen and stored at −80°C overnight. Total RNA was isolated using the Absolutely RNA miniprep Kit (Agilent) according to the manufacturer’s instructions. RNA quantification was measured by Nanodrop, and quality was assessed using the Prokaryote RNA Nano kit in an Agilent Bioanalyzer 2100 instrument. RNA was verified to be free from gDNA contamination using end-point PCR with primers listed in **Table S1**. Real-time PCR was performed using the iTaqUniversal SybrGreen one-step kit (BioRad, Hercules, Ca, USA) according to the manual protocol. Amplification of 300 ng input RNA was measured in an Eppendorf Mastercycler RealPlex thermocycler using the following parameters: 10 min at 50°C for the reverse transcription reaction followed by 1 min at 95°C to activate the DNA polymerase; Then 25 cycles amplification including 15 sec at 95°C for denaturation; 30 sec at 60°C for annealing; 1 min at 72°C for the extension step. One last step of 10 min was added for the final extension. The cycle threshold (C_t_) was automatically determined by the software (Eppendorf RealPlex 3.5). The melting curve was obtained after each run to ensure the specificity and quality of the products of each experiment. Experiments were performed in triplicate, and each sample was loaded in three technical replicates into each plate. Differential expression was calculated from the negative difference in threshold crossing point (-ΔC_t_) between the primers of interest and control primers that target a constitutively expressed housekeeping gene (*hvo_0136,* encoding translation elongation factor Eif-1A, **Table S1**).

### TF-DNA binding assays with chromatin immunoprecipitation coupled to quantitative PCR (ChIP-qPCR) or sequencing (ChIP-seq)

Cultures (50 mL) of HV284 (*troR-HA*), HV176 (*sirR-HA*), and HV177 (*idr-HA*) were grown in PR+ and PR-media up to OD of 0.3-0.5. Aliquots (45 ml) of each culture were analyzed by ChIP-seq as previously described (39) with modifications. In brief, DNA-protein crosslinks were generated by the addition of 1.4 mL of 37% (v/v) formaldehyde stock to the 45 mL culture aliquot to obtain a final concentration of 1% (v/v) formaldehyde. The samples were incubated in 200 ml pyrex flasks for 20 min at room temperature on a rocking platform (Thermo Scientific orbital shaker model 2314) to ensure homogeneous distribution of formaldehyde. To stop the crosslinking reaction, 5 ml of 1.25 M glycine was added to obtain a final concentration of 0.125 M. Cells were harvested by centrifugation (5,000 x *g*, 5 min at 4°C) in an Eppendorf centrifuge 5804R. Pellets were washed 3 times with 10 mL PR-and pelleted. Cells were lysed in 800 µL of lysis buffer (1 mM EDTA, 0.1% Na-deoxycholate, 140 mM NaCl, 50 mM HEPES-KOH pH 7.5, 1% Triton, 1x Halt^TM^ protease inhibitor cocktail 100x (Thermo Scientific #78429) and sonicated for 15 cycles of 1 min, 30 sec on/off each (Diagenode sonicator). Cell lysates were centrifuged (21,000 x *g*, 10 min at 4°C) to remove cell debris. Cell free extract (500 μl) was incubated with HA-antibody-bound beads (30 μl, equilibrated in PBS) overnight at 4°C in a rotating clamp. The next day beads were recovered using a magnetic platform, and washed thoroughly: twice with 1 ml of the previously described lysis buffer; twice with 1 ml of wash buffer (1mM EDTA, 10mMTris-HCl pH 8.0, 0.5% NP40, 250 mM LiCl and 0.5% Na-deoxycholate), and then once with TE buffer (1mM EDTA, 10mM Tris-HCl pH 8.0). 50 µl of elution buffer (50 mM Tris, 10 mM EDTA, 1% SDS (w/v), pH 8.0) was added to the beads 65°C for 10 min. Beads were separated from supernatant using the magnetic platform and 40 µl were used for IP cross-link reversal. 160 µl of TE/SDS was added to the samples and each tube was incubated at 65°C overnight to reverse cross-link. The samples were diluted with 1x volume of TE buffer (200 µl) and treated with RNase A. Finally, DNA was extracted by addition of 1x vol phenol: chloroform: isoamyl-alcohol (25:24:1) and purified with 1x vol 100% cold ethanol. DNA was washed with 500 µl of cold 80% ethanol and resuspended in 50 µL picopure water. Aliquots of the DNA sample (1 µl each) were used for DNA quantification by nanodrop and DNA quality analysis using High Sensitivity DNA Chips with an Agilent Bioanalyzer.

For ChIP-qPCR, immunoprecipitated DNA samples were analyzed using primers described in **Table S1** in technical triplicate using iTaq Universal SybrGreen reagents (BioRad, Hercules, Ca, USA) in a RealPlex Instrument (Eppendorf) with 30 sec Polymerase activation and DNA denaturation at 95°C, and 35 cycles with 15 sec denaturation at 95°C and 30 sec extension at 60°C. The melting curve was settled to increase the temperature from 65-90°C with 0.5°C increments of 2 to 5 sec per step. Enrichment for putative TroR and SirR binding sites was calculated by normalizing binding site qPCR product C_t_ values to C_t_’s from the 3’ end of the coding sequence of the gene of interest and input DNA background as previously described (40).

For ChIP-seq, DNA samples were quantified using the fluorometric quantitation Qubit 2.0 system (ThermoFisher Scientific). ChIP-seq libraries were prepared by Duke University Sequencing and Genomic Technologies Core Facility (Duke Sequencing Facility) from 1 ng of ChIP DNA using the KAPA HyperPrep Library Kit (Roche) to generate Illumina-compatible libraries. During adapter ligation, dual unique indexes were added to each sample to enable multiplexing. The resulting libraries were cleaned using SPRI beads, quantified, and checked for quality using Qubit 2.0 and Agilent Bioanalyzer. Libraries were pooled into equimolar concentration and sequenced by the Duke Sequencing Facility on an Illumina HiSeq 4000 sequencer with 50 bp single-end reads. Analysis was performed as described in (41). In summary, FastQC software (http://www.bioinformatics.babraham.ac.uk/projects/) was used to determine the quality, length, and GC distribution, as well as the proportion of adapters. Adapters were removed from the raw sequences using TrimGalore! (http://www.bioinformatics.babraham.ac.uk/projects/). The resultant sequences were aligned to the *Hfx. volcanii* genome (DS2, NCBI RefSeq GCF_000025685.1_ASM2568v1) using Bowtie2 (42), generating a SAM file for each sequenced sample. These files were compressed to binary and sorted to generate sorted BAM files. These BAM files were used as input for peak calling software. Binding peaks for the TF were called using the R Bioconductor package MOSAiCS (43) with the default parameters (two signal, FDR = 0.05, minimum peak size = 50). Peaks were assigned to genes and intergenic sequences within 500 bp of the peak center using code from (41). Genes were excluded for further analysis if the corresponding peak was located discordant with the direction of transcription. The peak neighboring *hvo_0531* was removed, as proteins binding this region are known to cross-react with the HA epitope (44). SirR ChIP-seq peaks were subjected to additional filtering steps due to technical variations in the input control read depth: peaks with average MOSAiCS score < 1000 were removed, peaks within 500 bp of genes annotated as transposases or other mobile elements were removed by manual curation. For all datasets, peaks were visualized and verified manually using the Javascript version of the Integrated Genome Viewer (IGV) (45). Genome-wide and zoomed regions for manuscript figures were generated using the R Bioconductor package trackViewer (46). SirR and TroR ChIP-seq analyzed data are provided in **Tables S2** and **S3**, respectively. Raw and analyzed data are accessible via Gene Expression Omnibus (GEO) at accession GSE240622.

### Western blot

Cell lysates diluted in commercial 2X Laemmli buffer (BioRad, Hercules, Ca, USA) were heated 3’ at 95°C, subject to SDS-PAGE (Mini-Protean precast gels, Any KD^TM^ BioRad, Hercules, Ca, USA) in 1X Tris-glycine-SDS running buffer, and transferred to nitrocellulose membranes (iBlot Transfer Stack, Invitrogen, Thermo Fisher Scientific). Blocking, wash, and antibody incubation steps were with rocking. Membranes were blocked with 25 ml 5% milk in TBST (20 mM TrisHCl, pH 7.5; 150 mM NaCl, 0.1% Tween20) for 1h at room temperature, washed 3x with 15 ml TBST for 10’. After blocking, the primary antibody (anti-HA, Abcam, ab9110) was added in 1:10,000 dilution and incubated at 4°C overnight. Membranes were subsequently washed 3 x 10 min in 15 ml of TBST, incubated with HRP anti-rabbit secondary antibody (Abcam, ab97051) at 1:50,000 dilution in TBST for 1h at room temperature. Bands were visualized using electrochemiluminescence Pierce ECL substrate (Thermo Scientific, protocol as suggested by the company), and Fujifilm were used for visualization.

### RNA-seq experiments and analysis

Total RNA was extracted from 50 mL cultures grown in PR+ and PR-media as described above. To deplete ribosomal RNA, 3-5 µg of each sample was treated with Ribozero® rRNA Removal Kit according to the manufacturer’s instructions (Illumina, USA, now discontinued. See reference (47) for updated rRNA removal protocols for haloarchaea). The resultant mRNA was precipitated with 100% cold ethanol and resuspended in 10 µL picopure water. Samples were verified to be free of rRNA in a second round of Agilent Bioanalyzer analysis. Libraries were constructed using ∼50 ng of mRNA using the Kapa Stranded RNA-Seq Library Preparation Kit (KAPA Biosystems, Illumina). Indexes were designed according to TruSeq RNA UD specifications (#20020591) and obtained from Integrated DNA Technologies. Resultant libraries were tested for quality and quantity by Bioanalyzer using the 2100_expert_High Sensitivity DNA Assay chip (Agilent Technologies) and sequenced on a Hiseq4000 instrument by the Duke Sequencing Facility. Results were quality checked, and reads were mapped to the *Hfx. volcanii* genome as described above and previously (41). R package DESeq2 was used to detect differentially expressed genes (48). In Δ*troR* vs Δ*pyrE* comparisons, resultant gene expression is given as a ratio of Δ*troR* : Δ*pyrE* in iron replete conditions. For iron response in the parental strain, gene expression is given as a ratio of Δ*pyrE* in the absence of iron relative to Δ*pyrE* grown in the presence of iron. Significant differential expression was defined as log2 fold change (LFC) ratio ³ |1| and adjusted *p*-value £ 0.05.

### NanoString gene expression experiments and analysis

Cell culture of parent (Δ*pyrE*) and Δ*arsR* mutant strains were grown in PR+ or PR-until mid-log phase (OD_600_ = 0.4-0.6); RNA from three biological replicates each strain and each condition was extracted and verified to be clean from DNA contamination and then quality tested by Bioanalyzer as described above. Nanostring gene expression assays were performed by the Duke Microbiome Center using the NanoString nCounter MAX Analysis System (Seattle, WA) (49). Customized probes were designed specific for detecting each of 30 TroR target genes and 3 housekeeping gene internal controls: *hvo_0717* (*ftsZ1*)(50), *hvo_0136* (*eiF1*α), and *hvo_2433* (*cox15*). RNA (100 ng) for each sample was hybridized to the probes in the assay, and the raw counts were obtained by the Duke Microbiome Core using nCounter software (NanoString, Seattle, WA). Raw counts were first background subtracted by thresholding to the geometric mean of Nanostring negative control probes (any probes detecting gene expression below this value were set to a floor of 1 count). Thresholded values were normalized to the geometric mean of the Nanostring-specified positive control probes and the *Hfx. volcanii* housekeeping genes. Log2 fold change ratios of Δ*arsR*: Δ*pyrE* expression from resultant normalized count data were calculated for each of PR-and PR+ conditions. *P*-values of significance vs the expected mean of 0 for each gene were calculated by 1-sided Student’s t-tests. Genes were considered significantly differentially expressed in the Δ*arsR* background relative to the parent strain if they met the criteria of: (a) *p*-value £ 0.05; (b) LFC ratio ≥ |1|. Nanostring probe sequences, raw data, normalized data, ratios, and statistics are listed in **Table S4**.

### Motif detection

For motif discovery, MEME version 5.5.1 was used within the MEME suite of online tools (51). DNA sequences corresponding to the peaks 140-210 nt in length from ChIP-seq data in the 5’ direction, were used as input. Default parameters were used to run MEME, except that motif width was constrained to 6-20 nucleotides. The MEME motif output (motif with lowest p-value) was used as input to FIMO genome-wide analysis (MEME suite) to find individual occurrences along a set of 4,023 DNA sequences upstream of gene coding sequences from the genome database *Hfx. volcanii* GCF 0000025685.1 ASM2568v1 assembly, February 23, 2023. Upstream was defined as 1,000 nt upstream to 200 nt within the coding sequences. A default threshold that includes motif sites in the output with a *p*-value ≤ 1.0 x 10^-4^ was used.

### Transcriptional reporter plasmid construction and assay

The promoter of *hvo_B0047* was amplified by PCR using Taq DNA polymerase as recommended by the company [New England Biolabs (NEB)]. The PCR primers flanking the region of interest were designed to include XbaI and NdeI sites in the corresponding forward and reverse oligos (**Table S1**). The plasmid pJAM2678 containing the *bgaH* gene encoding the β-galactosidase enzyme from *Haloferax alicantei* (52) was linearized using XbaI and NdeI compatible restriction enzymes (NEB) and purified from TAE agarose gels as explained previously (53). The PCR fragments of interest were treated with the same enzymes and cleaned by PCR cleanup (NEB). Linear plasmid pJAM2678 and each PCR product were ligated together in a reaction using T4 DNA ligase overnight at 16 °C to obtain plasmid pAKS202 that contained 150 bp upstream of the translation start site of *hvo_B0047* cloned in front of *bgaH*. Plasmids pAKS204, pAKS205, and pAKS207 (**Table S1**) were generated by PCR based on the Q5 site-directed mutagenesis kit, where one or both motifs have been degenerated using the primers designed with the NEB online primer design software NEBaseChanger.neb.com (primers listed in **Table S1**). The parental (Δ*pyrE*) and Δ*troR* strains were transformed with: (a) pJAM2678 positive control plasmid carrying a strong constitutive promoter P2 (53); (b) pJAM2714 negative control plasmid carrying only the Shine-Dalgarno ribosome binding sequence of ribosomal gene P*2rrnA* (53); (c) pAKS202; (d) pAKS204; and (e) pAKS207. Three independent colonies of all constructions were grown as previously described in PR+ and PR-media, and the levels of β-galactosidase activity were quantified as explained in (54). Briefly, cell lysates containing equimolar amounts of protein estimated by Bradford assay (55) from the different strains grown in the presence or the absence of iron were added into the ONPG buffer [2.5 M NaCl, 10 µM MnCl_2_, 0.1% (w/v) β-mercaeptoethanol, 50 mM Tris-HCl, 1.33 mM *o*-nitrophenyl-ß-D-glucopyranoside (from stock prepared with 8 mg.mL^-1^ ONPG in 100 mM potassium phosphate buffer, pH 7.2)]. Absorbance was measured at 405 nm. One unit of enzymatic activity corresponds to the amount of the enzyme catalyzing the hydrolysis of 1 µmol of ONPG per min. Ratios of enzymatic activity in iron replete vs iron poor conditions are reported in the figures and text. Statistical significance was determined using two-tailed, two-sample homoscedastic Student’s *t*-tests. Resultant *p*-values are reported in the text.

## RESULTS

### DtxR transcription factor family is extensively and specifically expanded in Haloarchaea

To determine the phylogenetic conservation of the DtxR family, all available archaeal genome sequences in the database were queried across phyla (see Methods). DtxR orthologs were detected in 1,299 queried genomes (**Fig 1A**). DtxR orthologs are more common in euryarchaeal lineages: the number of euryarchaeal orthologs include 160 in the Methanobacteria, 284 in the Thermoproteota (formerly Marine Group II archaea), and 516 in the Haloarchaea. Of these euryarchaeal lineages, the highest percentage of genomes containing DtxR orthologs is found in the Haloarchaea (39.72%). In addition, a swift recent radiation greatly expanded the number of DtxR family orthologs within Haloarchaea (**Fig 1A**). In the current query, 304 haloarchaeal genomes included each contained 1-6 DtxR orthologs. This finding expands the number from a previous study in which each of 11 haloarchaeal genomes analyzed encoded 2 to 4 paralogs of this TF family (19). The *Hfx. volcanii* genome encodes three predicted DtxR metalloregulators: TroR (HVO_0863), Idr (HVO_0538), and SirR (HVO_0819). All three contain the expected DtxR domains based on homology with bacterial DtxR homologs (56). These domains include an N-terminal helix-turn-helix (HTH) DNA-binding domain and a C-terminal dimerization domain with two metal binding sites (**Fig 1B**, top). One of these DtxR proteins, SirR, also contains an SH3-like domain as an extra C-terminal extension (**Fig 1B**). These three *Hfx. volcanii* paralogs exhibit variable levels of sequence conservation to one another across the protein, with TroR and SirR exhibiting the highest protein sequence identity (33%, **Fig 1B**, bottom). A multiple sequence alignment between these paralogs suggests that they share higher sequence identity in the N-terminal HTH domain than in the C-terminal metal binding domain (**Fig 1B** and **C**). Taken together, these data suggest that: (a) the DtxR family of TFs is widely conserved across archaea but specifically expanded in the Haloarchaea; (b) the model haloarchaeal species *Hfx. volcanii* encodes three canonical DtxR family paralogs that are strong candidates for further study of iron regulation in archaea.

### *troR* is an essential gene whose product is necessary for growth under variable iron conditions

To determine the iron conditions that support the growth of *Hfx. volcanii*, growth of the parent was examined in minimal media under a wide range of iron concentrations under aerobic conditions. Cultures grew poorly in iron-depleted media (**Fig 2A**), with an average maximal growth rate of ∼0.06 h^-1^ (**Fig 2B**). These iron starvation growth rates were statistically indistinguishable following one (1x noFe) or two passages (2x noFe) in iron-poor medium, where a “passage” is defined as overnight incubation in a medium treated with the iron-chelating agent BPS (see Methods). In contrast, iron supplementation improved growth (**Fig 2A**), reaching ∼2 times faster rates in 100 µM FeSO_4_ (∼0.11 h^-1^). This concentration of iron supports significantly faster growth than 10 and 250 µM FeSO_4_ (BH corrected pairwise Student’s t-test *p* < 0.013 and < 1.52 x 10^-4^, respectively, **Figure 2B**). Together these results suggest that one passage in the iron-poor medium is sufficient to impair growth, and 100 µM FeSO_4_ supplementation in a minimal medium supports optimal growth of *Hfx. volcanii*. These conditions of iron [PR-(no iron is added) and PR+ (100 µM FeSO_4_ is added) constitute the “depleted” and “replete” conditions, respectively] were therefore selected for subsequent experiments.

**Figure 2.**
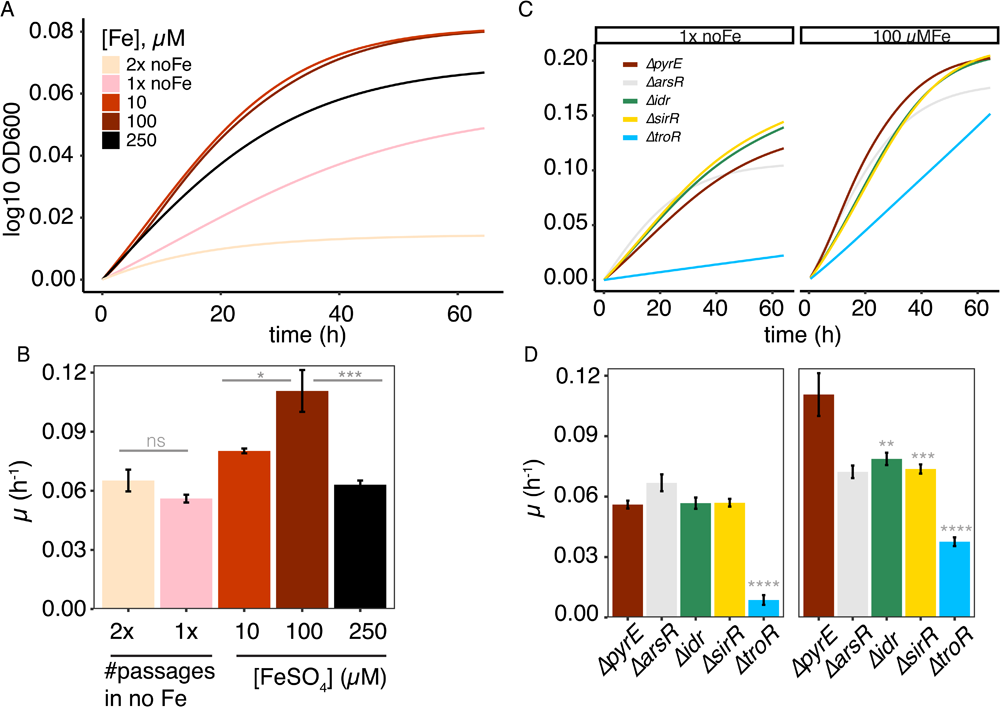
*Hfx. volcanii* TroR is required for growth in varying iron concentrations. (A) Average growth curves (logistic regression fit to the log10-transformed raw data) of the Δ*pyrE* parent strain H26 using the growthcurver package in R (26). Colors corresponding to each iron condition are shown in the legend, and colors are consistent across panels A and B. 2x noFe, cells were grown to saturation twice in minimal medium with no added iron (PR-, see Methods), with washing in between passages. 1x noFe, cells grown were grown to saturation once in PR-medium to before the growth curve shown is shown. The number of replicate curves comprising each condition are as follows: 2x noFe, n = 9; 1x no Fe, n = 18; 10 µM Fe, n = 9; 100 µM Fe, n = 18; 250 µM Fe, n = 9. (B) Maximum growth rate (µ) in hours^-1^ across curves shown in A, with each bar representing the mean of the number of replicates listed above, and error bars the standard deviation from the mean. Asterisks represent significant *p*-values from two-sided Student’s *t*-tests with Benjamini-Hochberg (BH) correction for multiple hypothesis testing. \**p* < 0.05; \*\**p* < 0.01; *** *p* < 0.001; **** *p* < 1 x 10^-04^. (C) Average growth curves of the Δ*pyrE* strain H26 compared to the four TF mutant strains of interest. Colors corresponding to each strain are given in the legend. The number of replicate curves comprising each condition are as follows: Δ*arsR*, n = 3; Δ*idr*, n = 12; Δ*sirR*, n = 12; Δ*troR*, n = 12; Δ*pyrE*, n = 18. (D) Maximum growth rate (µ) in hours^-1^ across curves shown in C, with each bar representing the mean of the number of replicates for each strain listed in C, and error bars the standard deviation from the mean. Asterisks result from the same type of tests as in B -- comparing test *vs* control -- and have the same designations as in (B).

To better understand the role of the three *Hfx. volcanii* DtxR paralogs in maintaining cellular physiology in response to iron, strains deleted for each DtxR-encoding gene were constructed (Δ*troR*, Δ*idr*, and Δ*sirR*). To validate the genotype of these mutants, whole-genome sequencing analysis was performed allowing the detection of potential secondary mutations as well as any wild type copies of the gene targets that may remain in the polyploid genome (41,57,58). Each strain appeared to represent an in-frame deletion by PCR and Sanger sequencing, and genome resequencing verified that no remaining genomic copies of the *troR*, *idr*, and *sirR* genes were detected in the corresponding mutant strains (**Fig S1**). In addition, Δ*idr* and Δ*sirR* were free of second-site mutations within protein coding sequences, suggesting that any phenotype observed would be a consequence of the deletion of the gene of interest (**Table S1**). However, a second-site C to T mutation in position 1,633,902 of the main chromosome (NCBI RefSeq ID NC_013967.1 (32)) was detected in the Δ*troR* strain (**Table S1**). This mutation changes codon 48 [CGC, arginine (R); to TGC, cysteine (C)] in the DNA binding domain of HVO_1766 (WP_004041648.1), an ortholog of a bacterial ArsR family transcriptional regulator. The first cytosine residue from the trio R48 is conserved across bacteria and is considered to be part of the winged helix-turn-helix domain responsible for the DNA binding, so mutation of this nucleotide is predicted to abolish ArsR-DNA binding (59). Bacterial ArsR family members have been implicated in regulation of a variety of metal and metalloid responses (59,60), so an Δ*arsR* (Δ*hvo_1766*) deletion strain was generated as an important control for the metal response of the Δ*troR* deletion strain in subsequent experiments. This Δ*arsR* strain was verified by whole genome sequencing to be free of all genomic copies and second site mutations (**Fig S1** and **Table S1**). To further test for *troR* essentiality, we attempted to re-construct the Δ*troR* deletion while ectopically expressing wild type *arsR in trans* (pAKS190, **Table S1**). However, three trial transformations yielded no viable colonies. Δ*troR* was able to be transformed with empty vector negative control, indicating that neither antibiotic selection nor the pAKS190 plasmid itself were the cause of the toxicity. As an additional control, we successfully transformed pAKS190 in the Δ*arsR* strain, showing that transformation with this plasmid is feasible. Together these experiments suggest that *troR* is essential for viability, and that mutation of *arsR* suppresses that essentiality. Therefore, we proceeded with the originally isolated Δ*troR* clone containing the *arsR* point mutation (R48C), and hereafter, the strain referred to as “Δ*troR”* contains this *arsR* point mutation.

To determine the physiological role of DtxR paralogs and ArsR in the iron response, the growth of all four mutant strains was compared to the parent strain in response to +Fe and -Fe conditions. The Δ*troR* mutant exhibited the strongest growth defect relative to all other strains investigated (**Fig 2C**). Growth of this mutant strain was barely detectable in iron poor conditions and reached only 34% of the parent growth rate in iron replete conditions (*p* < 1 x 10^-04^, **Fig 2D**). This slow growth was not due to the *arsR* point mutation in the Δ*troR* background because the Δ*arsR* mutant growth rate was only slightly slower and not significantly different from that of the parent (p = 0.0832 in PR-, 0.104 in PR+, **Fig 2D**). In contrast, the Δ*idr* and Δ*sirR* mutants showed a slight but significant growth defect relative to the parent strain under replete but not iron depleted conditions (*p* < 0.01 and < 0.001, respectively, **Fig 2D**). Taken together, these results suggest that Idr, SirR, and ArsR play minor roles in growth under PR-and PR+ conditions, but that *troR* is essential to cell viability.

### Analysis of the TRN of the DtxR TFs

To characterize the gene regulatory network (TRN) topology comprised by the DtxR TFs in *Hfx. volcanii*, we first asked whether the expression of genes encoding each TF is dependent on regulation by each of the other DtxR TFs. Thus, quantitative reverse transcriptase PCR (RT-qPCR) analysis was used to examine the relative transcript abundance of each of the TF-encoding genes in the presence and absence of iron in parent, Δ*troR*, Δ*idr*, and Δ*sirR* strains (**Fig 3A**). In the parent, the abundance of *idr* transcripts was reduced approximately 6-fold in the presence vs. absence of iron. Deletion of *sirR* did not perturb this response (**Fig 3A**, left panel). In contrast, the expression levels of *idr* transcripts remained elevated in the Δ*troR* strain in the presence of iron, suggesting that TroR is necessary to repress *idr* expression in this condition (**Fig 3A**, left). The expression levels of *sirR* and *troR* transcripts remained consistent across iron conditions and were unaffected by deletion of the other DtxR TFs (**Fig 3A**, middle and right panels). This result suggests that *sirR* and *troR* transcripts are constitutively expressed and are not regulated by other DtxR TFs.

**Figure 3.**
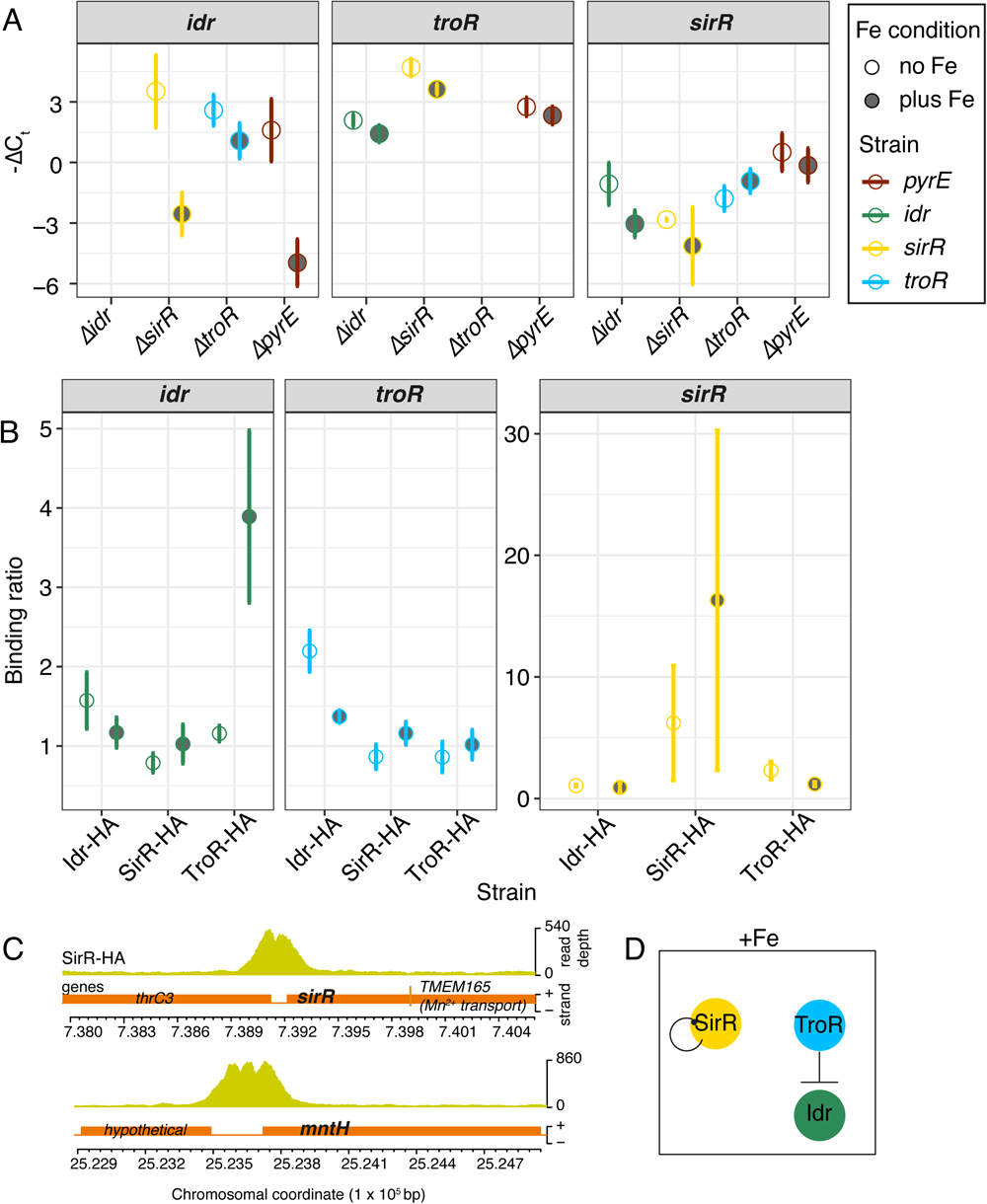
The Gene Regulatory Network of the DtxR proteins in *Hfx. volcanii* is governed by TroR in the presence of iron. (A) Relative gene expression of *idr, sirR*, and *troR*, measured by qRT-PCR in the parental strain (dark red), and in the three knockout strains Δ*idr* (green), Δ*sirR* (yellow), and Δ*troR* (blue) in the presence (filled dots) and absence of iron (empty dots). Values are shown as −ΔC_t_ relative to the housekeeping control gene HVO_0136. Error bars represent the standard deviation of three biological replicates and three technical replicates. (B) The binding ratio of the epitope-tagged DtxR proteins Idr:HA, SirR:HA, and TroR:HA to the promoters of genes *idr* (green)*, sirR* (yellow), and *troR* (blue), measured by coupling qPCR to chromatin immunoprecipitation (ChIP-qPCR). Primer sequences are included in ST1. Each dot represents the average of three biological replicate trials. Empty dots represent iron-poor conditions; filled dots iron-replete conditions. Error bars represent the standard error of the mean of the binding enrichment. (C) Visualization of SirR-HA ChIP-seq binding enrichment upstream region of the *sir* gene (HVO_0819) (top). Read pileups (Y-axis) for a representative biological replicate experiment are shown above in yellow, genes below in orange. Detailed gene annotations are provided in **Table S2**. Briefly, *thrC3, HVO_0818,* threonine synthase (74); hypothetical, gene of unknown function. The vertical orange bar represents a 3 bp overlap between the *sirR (HVO_0819)* 3’ end and the 5’ end of the downstream gene (*HVO_0820,* encoding TMEM165 family protein implicated in Mn^2+^ transport (68)). Shown below is a visualization of SirR-HA binding enrichment upstream of *mntH* (*HVO_2674*, encoding an N-ramp family Mn^2+^ transporter). In each zoomed visualization, X-axis represents the Genomic coordinates (in 1 x 10^5^ bp) of NC_013967.1 (main chromosome). All SirR peaks and their corresponding annotations are listed in **Table S2**. Peaks across all genomic elements and biological replicates are shown in **Figure S2**. (D) Schematic representation of the network topology of the DtxR family of TFs in *Hfx. volcanii* shows that in iron-replete conditions, SirR regulates its own expression and TroR regulates *idr* expression.

To investigate the binding of the DtxR TFs to the promoters of the genes encoding the other TFs, chromatin immunoprecipitation coupled with quantitative PCR (ChIP-qPCR, primers in **Table S1**) was performed. The hemaglutinin (HA) epitope was translationally fused to the C-terminus of each DtxR TF to produce the strains Idr-HA, SirR-HA, and TroR-HA. We observed strong enrichment of TroR-HA to the *idr* promoter only under iron-replete conditions (∼4x fold enrichment), consistent with RT-qPCR data and further suggesting that TroR represses *idr* in the presence of iron (**Fig 3A and B**). In contrast, no TroR-HA binding was detected at the promoters of *sirR* and *troR* (**Fig 3B**). SirR-HA binding was observed to be enriched at the *sirR* promoter, indicative of autoregulation, but was not detected upstream of *idr* or *troR* (**Fig 3B**, right-most panel). ChIP-qPCR enrichment at the *sirR* upstream region was highly variable across biological replicates. To validate SirR binding in the promoter of its own encoding gene, chromatin immunoprecipitation coupled with DNA sequencing (ChIP-seq) was performed. Such analysis revealed the existence of a strong peak of binding enrichment in the promoter of *sirR* in the SirR-HA (vs. parent) background, consistent with the hypothesis that *sirR* is autoregulated (**Fig 3C**). SirR-HA was found to bind to only 12 other sites in the genome (nearby genes within 500bp of the peak center located in the direction of transcription), primarily nearby genes of unknown function (**Table S2, Fig S2**). However, among these 12 sites, SirR-HA was found to bind 5’ of gene encoding an Nramp family manganese uptake transporter homolog (HVO_2674) (61,62). These data suggest that SirR is more likely to regulate manganese uptake and may not be centrally involved in the iron response. Together, these results suggest that the *Hfx. volcanii* DtxR TRN is comprised of three TF nodes and two edges (influence of expression regulation): TroR represses *idr* expression in the presence of iron, and SirR binds its own promoter under iron-replete conditions (**Fig 3D**). The *Hfx. volcanii* DtxR TRN architecture appears to be far simpler and less interconnected than that of *Hbt. salinarum* (9).

### TroR is required for repression of iron uptake and activation of iron storage

Because TroR plays a major role in response to iron imbalance (**Figs 2 and 3**), the genome-wide regulon (*i.e*., set of target genes) under the control of TroR was next characterized using functional genomic sequencing methods. ChIP-seq analysis was performed under iron replete conditions where TroR was expected to function as a regulator (**Figs 2 and 3**). TroR was observed to reproducibly bind 36 specific regions of the *Hfx. volcanii* genome (**Table S3** and **Fig S2**). These TroR binding “peaks” were located within 500 bp of 54 genes, with the majority occurring within non-coding regulatory regions (**Fig 4A**, **Table S3**).

**Figure 4.**
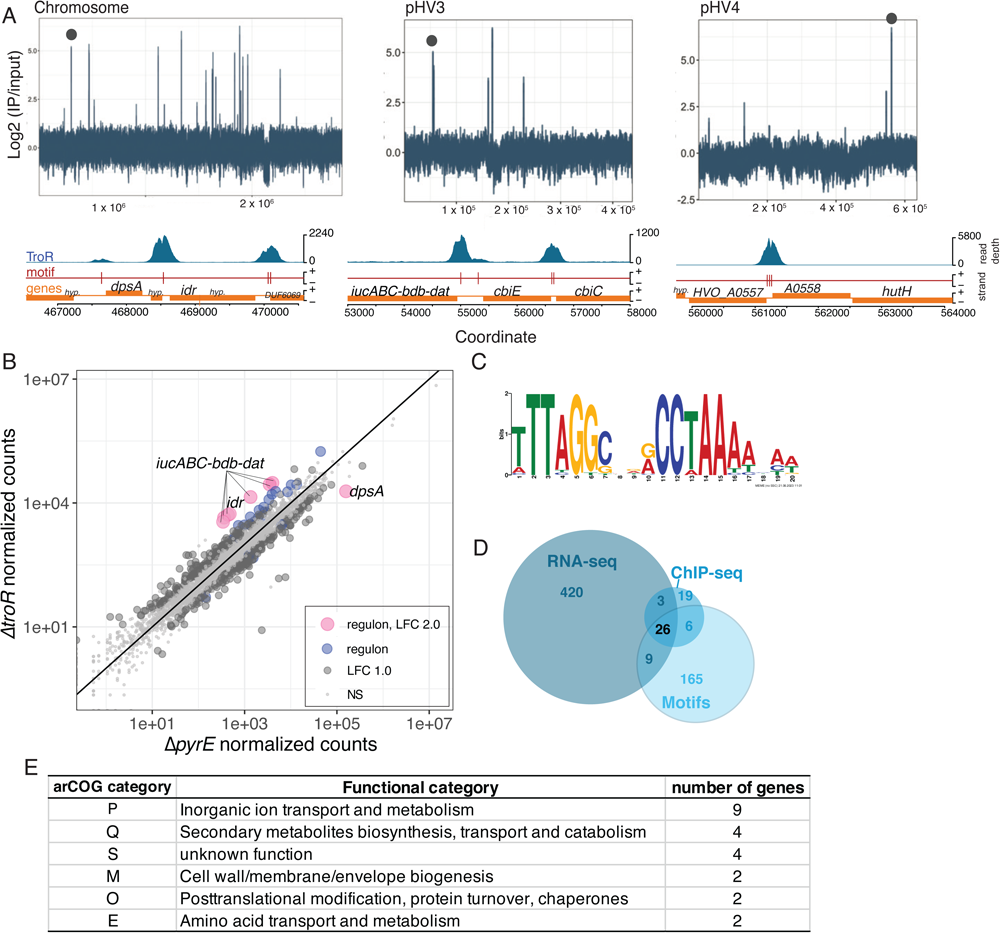
The TroR regulon governs iron-dependent expression of genes encoding functions in iron uptake and storage. (A) Visualization of TroR binding enrichment peaks across the genome. Upper panels show genome-wide binding enrichment (x-axis: ratio of IP enriched TF-DNA binding vs input control, y-axis genomic coordinate). Black dots are used to mark the position of the zoomed-in cis-regulatory TroR binding motifs and genes (from representative experiments) below each chromosomal element (from left to right, main chromosome, megaplasmid pHV3, pHV4). Detailed annotations and unique gene identifiers for each gene are given in **Table S3**. Briefly, *hyp*., hypothetical gene of unknown function; *dpsA*, iron storage; *idr,* metal-dependent transcriptional regulator; DUF, domain of unknown function; *iucABDC-bdb-dat*, siderophore biosynthesis gene cluster (20); *cbiCE,* cobalamin (vitamin B12) biosynthesis genes; *HVO_A0557-8,* Fe^3+^-hydroxamate ABC transporter; *hutH,* histidine ammonia lyase. All TroR peaks and their corresponding annotations are listed in **Table S3**. Peaks across all genomic elements and biological replicates are shown in **Figure S2**. (B) Quantitation of gene expression (RNA-seq normalized read counts) in Δ*troR* mutant (y-axis) vs Δ*pyrE* parent strain (x-axis) in the presence of iron. Pink points represent genes whose expression is strongly affected by deletion of *troR* and are part of the TroR high confidence regulon (bound by TroR, have a motif in their promoter, and are significantly differentially expressed by at least Log2Fold Change (LFC) LFC>2.0 and BH adjusted *p*-value £ 0.05). Blue points represent genes in the TroR regulon differentially expressed 1<LFC<2 and *p* £ 0.05. Dark grey, LFC>1.0 and *p* £ 0.05. Light grey, not significant (NS). (C) Cis-regulatory binding motif for TroR detected within ChIP-seq binding peaks. Y-axis represents motif bit scores, x-axis the position of each nucleotide in the motif in bp. (D) Venn diagram depicting the number of genes detected within the intersection of each dataset. 26 genes are observed in all three datasets and are therefore members of the high confidence TroR regulon (see also text). (E) arCOG gene ontology functional categories of the genes in the high confidence regulon of TroR. See Table S3 for additional detail on gene annotations. Categories with fewer than 2 genes were not included.

To determine whether these TroR binding events resulted in significant differential expression, genome-wide transcriptome analysis (RNA-seq) was conducted. The datasets were first queried for genes that were significantly differentially expressed at the transcript level in response to iron availability in the parent (**Table S3**). Of the 3,870 genes for which expression was detected in RNA-seq experiments, 26% were significantly upregulated in the absence vs presence of iron [log2 fold change (LFC) ≥ |1|, p-value ≤ 0.05], whereas 23% were significantly downregulated (**Table S3**). Therefore, iron availability is associated with large-scale genome-wide changes in expression in the parent. When examining transcript abundance in the presence of iron, 458 genes were significantly up-or down-regulated in the Δ*troR* strain compared to the parent (Δ*troR* : parent LFC ≥ |1| and *p*-value ≤ 0.05, **Fig 4B, Table S3**). Of these, 20 genes were also bound by TroR in their upstream region according to ChIP-seq, suggesting that TroR is responsible for their iron-dependent regulation. 15 of these 20 genes that neighbored ChIP-seq peaks were members of operons (a cluster of genes that might be regulated together (63)). Genes that were members of these operons, downstream of the ChIP-seq peak (in the correct transcriptional direction), and differentially expressed in Δ*troR* were also considered to be regulated by TroR, bringing the number of genes potentially regulated by TroR to 29 (**Fig 4B and D**).

To provide further evidence that TroR regulates these genes, *de novo* computational cis-regulatory binding motif discovery was conducted using intergenic DNA sequences bound by TroR in ChIP-seq experiments as input (32 of 36 total detected peaks, Methods). The entire *Hfx. volcanii* genome was then scanned for this motif. A semi-palindromic motif TTTAGGC-NNN-CCTAAAANAN was detected in 206 instances in this genome-wide scan (**Fig 4C and D**). This motif was detected in all 32 peaks, some of which were located within non-coding regions upstream of large operons (see below). This provides strong validation that the ChIP-seq experiment captured true TroR binding events (**Fig 4A, C, and D**). Many of the TroR binding sites contained more than one instance of the motif and, in most cases, these multiples were detected in the promoter regions of genes involved in ion transport and metabolism (*e.g.*, siderophore biosynthesis and iron ABC transporters, **Fig4A** and **Table S3**).

Integrating the ChIP-seq, RNA-seq and motif discovery data together; 26 genes were commonly detected by all three methods, strongly suggesting that TroR directly regulates these genes by binding to its specific cis-regulatory motif in an iron-dependent manner. This list of 26 genes is named the “high confidence regulon of TroR” (**Fig4D** and **E**). As a control, we tested whether any of these genes were also regulated by ArsR using gene expression analysis via Nanostring mRNA counting technology (21,49). Gene expression in Δ*arsR* vs the parent was compared in iron poor vs iron replete conditions. The 30 genes that were differentially expressed in Δ*troR* in RNA-seq experiments were tested (**Fig 4B**), 11 of which were members of the high confidence regulon (**Fig 4E, Table S3**). None of these genes were significantly differentially expressed in Δ*arsR* in any conditions tested (**Table S4**). These Nanostring data suggest that TroR controls its high confidence regulon independently of ArsR. Therefore, taken together these functional genomics data suggest that TroR directly controls 26 genes in the presence of iron.

When examining these TroR regulon genes by archaeal Clusters of Orthologous Genes (arCOG) gene functional ontology analysis (64), the majority of the encoded proteins are associated with ion transport and metabolism. Specifically, half belong to arCOG groups P and Q (**Fig 4E**), suggesting that the majority of TroR target genes function in transporting inorganic ions, including many that are annotated as iron and siderophore transporters (**Fig4A** and **E, Table S3**). Eight of these genes in the high confidence regulon were strongly dependent on TroR regulation (LFC > |3| and p-value < 0.01; **Fig 4A & B)**. Seven of these eight genes were significantly upregulated in Δ*troR* in the presence of iron, suggesting that they require TroR for repression under these conditions. Six of these seven are members of a large operon previously shown to encode siderophore biosynthesis and transport proteins (*iucABDC-dat-bdb*) (14,20), and one encodes the DtxR family TF Idr (HVO_0538), validating our results from the ChIP-qPCR and RT-qPCR analysis (**Fig 3**). The eighth gene, *dpsA,* was strongly downregulated in Δ*troR* and bound by TroR in iron replete conditions. *dpsA* encodes a protein essential for iron storage (16). These data suggest that TroR is necessary to activate expression of *dpsA* to enable iron storage during replete conditions. In summary, these results suggest that TroR regulates the transcription of a specific set of genes highly enriched for functions in iron uptake and storage.

### TroR directly regulates the siderophore biosynthesis gene cluster in an iron-dependent manner via a semi-palindromic cis-regulatory binding motif

To further understand the functional importance of the computationally detected motif in the regulation of the current TroR targets, gene expression validation experiments and promoter-reporter transcriptional fusion assays were performed. To validate the observation that TroR is required for repression of the siderophore biosynthesis gene cluster (*hvo_b0041-hvo_b0047*, *iucABDC-bdb-dat*, **Fig4A**), the expression of *hvo_b0046* (*dat*) encoding a predicted diaminobutarate-2-oxoglutarate transaminase involved in siderophore biosynthesis (14,20) was tested by RT-qPCR (**Fig5A**). In the parent, Δ*sirR*, and Δ*idr* strains, *hvo_B0046* expression was strongly induced in the absence of iron relative to iron replete conditions, as expected from the RNA-seq experiments (**Fig5A, Fig4B**). By contrast, *hvo_b0046* expression remained elevated regardless of condition in Δ*troR*, validating that TroR is required to repress transcription of this gene in the presence of iron (**Fig5A**).

To better understand how the TroR cis-regulatory binding motif (**Fig5B**) affects transcriptional activity at target genes, we fused the wild type motif 5’ of the siderophore biosynthesis operon (*iucABDC-bdb-dat,* promoter P*_hvo_b0047_*) to the gene encoding β-galactosidase (β-gal) (*bgaH* from *Haloferax alicantei* (52), pAKS202). We then constructed three disrupted versions of this motif and measured the resultant enzymatic activity of β-gal as a proxy for gene expression; the enzymatic activity was calculated as a ratio of iron deplete to iron replete conditions (**Fig 5C**).

**Figure 5.**
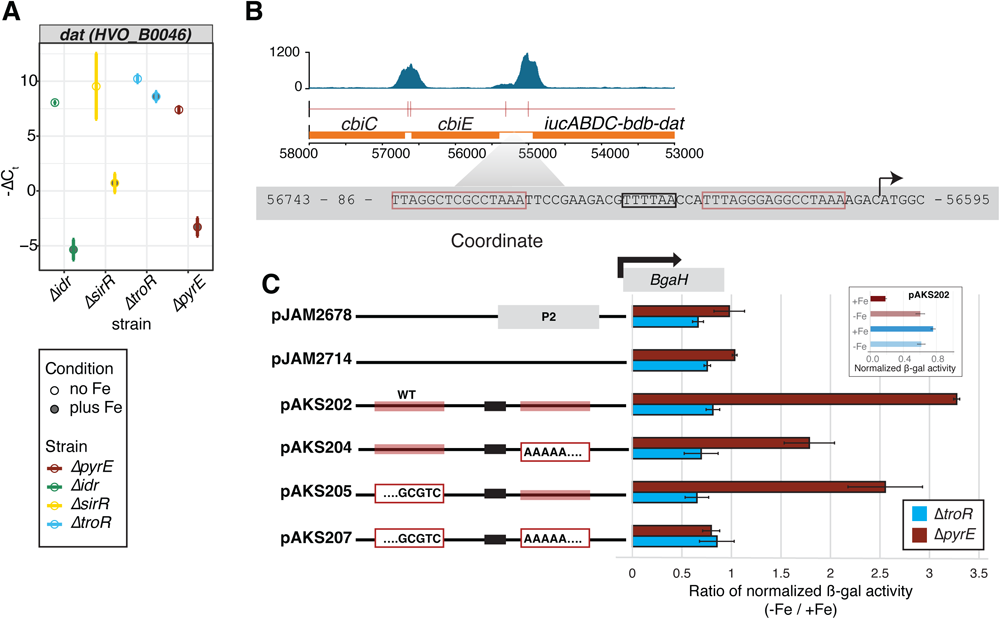
Examination of the TroR cis-regulatory motif and target gene expression. (A) Relative gene expression of the TroR target gene *hvo_b0046* measured by qRT-PCR in the parental strain (orange), and in the three knockout strains Δ*idr* (green), Δ*sirR* (yellow), and Δ*troR* (blue) in the presence (filled dots) and absence of iron (empty dots). Values are shown as −ΔCt relative to the housekeeping control gene *hvo_0136*. Error bars represent the standard deviation of three biological replicates and three technical replicates. (B) Zoomed region (top) corresponds to the siderophore biosynthesis gene locus, including the ChIP-seq peak, motif, and genes surrounding the promoter (P*_hvo_B0047_*) of interest. In the lower panel, the promoter sequence, including the predicted TATA binding site (black box) and TroR binding motif sequences (red boxes) are indicated. (C) At left, schematic representation of genetic constructions to test the activity of the P*_hvo_b0047_* promoter (see also **Table S1**). pJAM2678 is the positive control containing strong promoter P2. pJAM2714 is the negative control that lacks the TATA and TroR motif sequences. For constructs containing TroR binding motifs, wild type sequences are indicated by the red boxes, and TATA sequence are indicated by black boxes. Nucleotide substitutions by site-directed mutagenesis are indicated in white boxes with red borders. All promoter constructs are transcriptionally fused to the *bgaH* gene indicated at top. At right, the bar graph depicts the ratio of β-galactosidase enzymatic activity in iron poor (-Fe) divided by that in iron replete (+Fe) conditions. Ratios in the parent (dark orange bars) and Δ*troR* (blue bars) backgrounds are shown across promoter-reporter fusion constructs. Error bars represent the standard error of the mean of three biological replicate experiments. Upper box shows the normalized β-gal activity for the pAKS202 construct carrying the two unedited conserved motifs in iron replete (dark) and iron deplete conditions (light) in the parent strain (brown) and the Δ*troR* strain (blue).

Relative to controls, we observed that P*_hvo_b0047_* activity is significantly repressed 2-to 3.5-fold in iron replete vs. deplete conditions in the parent regardless of which of the two cis-regulatory binding motifs were present (pAKS202, 204, and 205, **Fig 5C**). Iron-dependent repression was lost when both regulatory motif sequences were mutated in the same construct (pAKS207). In contrast, promoter activity ratios remain constant across all constructs in Δ*troR* (**Fig 5C**). This finding is because promoter activity was constitutively high in the Δ*troR* background across constructs (see pAKS202 example, **Fig 5C** inset). Indeed, pAKS202 promoter transcriptional activity was significantly higher in Δ*troR* in iron-replete conditions relative to that of the parent strain (inset, *p* < 4.65 x 10^-5^). Taken together, these data support the functional genomics results (**Fig 4**) and suggest that: (a) TroR is required to bind at least one of the two cis-regulatory motif sites in P*_HVO_B0047_* to repress gene expression in the presence of iron; (b) the presence of either one of the two TroR binding motifs is sufficient for TroR-mediated repression.

## DISCUSSION

Here we show that the three DtxR family transcription factors (TFs) encoded in the *Hfx. volcanii* genome form a simple transcriptional regulatory network (TRN) in which TroR represses Idr under iron replete conditions, and SirR is autoregulated (**Fig 3**). In iron-replete conditions, TroR binds to a semi-palindromic DNA sequence motif to repress the promoters of 26 genes involved in iron uptake (siderophore biosynthesis genes) and activate iron storage gene expression (*dpsA*, **Fig 4** and **5**).

The *troR* gene appears to be essential for viability, and its expression is constitutive (**Fig 2** and **3**). Attempted deletion of *troR* selects for second site mutations in a gene encoding another TF, ArsR (HVO_1766). While the function of ArsR remains unclear in *Hfx. volcanii*, TFs of the ArsR/SmtB family in bacteria are classically known to repress genes encoding metal or metalloid (As(III), Zn(II), etc) efflux transporters using the metal of interest as an inducer (65). However, organisms that encode multiple ArsR family paralogs appear to have undergone functional divergence (66) such that a myriad of functions, including those other than metal homeostasis, have now been identified for bacterial ArsR family orthologs (59). Although no ArsR family homologs have yet been characterized in archaea, many halophilic archaeal genomes, including *Hfx. volcanii*, encode multiple ArsR paralogs (32). Interestingly, TroR binds to the ArsR promoter region in our ChIP-seq experiments (Table S3); however, no differential expression of *arsR* was detected in the Δ*troR* deletion strain, suggesting that TroR may not directly regulate *arsR* transcription. In further support of the idea that the genetic link between *troR* and *arsR* is likely indirect (perhaps to ameliorate toxic effects of Fenton reactions), we did not detect any co-regulation of TroR target genes here (Fig S3). Nevertheless, ArsR may play an important role in the metal homeostasis TRN of *Hfx. volcanii* and is worthy of future study.

SirR (HVO_0819) also likely plays a role in metal homeostasis, but our data suggest that it functions as a repressor of manganese uptake, consistent with its bacterial homologs (67). The *sirR* gene is likely expressed in operon with a TMEM family transporter given the 3 bp overlap in the coding sequences of these genes. The TMEM family has been implicated in Mn^2+^ transport in eukaryotes and bacteria (68). Therefore, SirR likely regulates this transporter by virtue of autoregulation of the *sirR* locus (**Fig 3C**). We also demonstrate that SirR binds to the promoter region of *mntH* (**Fig 3C**), encoding a homolog of Nramp family transporters which transport Mn^2+^ in bacteria (69). MntH has also been characterized by genetic analysis in *Hbt. salinarum* (61), to which the *Hfx. volcanii* homolog shares 64.84% amino acid sequence identity.

More broadly, the archaeal DtxR family of transcription factors (TFs) is strongly conserved with that in bacteria (19). In gram positive bacteria, DtxR homologs typically bind and repress genes encoding metal import (and toxin expression in pathogenic species) by binding their cognate metal ions (70,71). Each DtxR TF paralog within a given species appears to be dedicated to regulation of the uptake of a particular metal such as manganese, iron, zinc, etc. (72). In halophilic archaea, the number of DtxR members varies from one to six paralogs per genome across hypersaline adapted archaeal species (**Fig 1**). How DtxR TFs regulate metal homeostasis may therefore vary across species. For example, we showed previously that the DtxR metalloregulatory network in *Hbt. salinarum* is comprised of four interacting TFs that regulate each other (9). This cross-regulation between DtxR TFs forms an extensive network in which feedback loops enable dynamic tuning of gene expression in response to variability in iron concentrations over time (9). In addition, cross-regulation enables a broader homeostatic range of intracellular iron tolerance compared with bacteria (9). In contrast, we show here that *Hfx. volcanii* network is far simpler, with only three TFs and only one interconnection in which TroR represses Idr.

Although it remains unclear how the environment selects for divergence in TRN topology, based on the presented differences between *Hfx. volcanii* and *Hbt. salinarum* DtxR TRNs, we hypothesize that the complexity of the regulatory network might scale with the variability of metal bioavailability in the natural environment. In this regard, is interesting to observe that *Hbt. salinarum,* with a higher number of DtxR paralogs, colonizes pelagic niches of salt lakes and it has also been detected blooming in sporadic rainy seasons in the Dead Sea, when salinity of the water surface abruptly changed as a consequence of uncommon precipitation (73). Meanwhile, *Hfx. volcanii* colonizes the sediment of the Dead Sea (27), where metals in precipitation are plentiful, soluble ferric iron has also been detected and conditions are typically stable (29). Since Rodionov and Leyn (19) have predicted DtxR regulons across all halophiles, and differences on the number of DtxR paralogs have been found in each species, it will be interesting in the future to determine how these TRNs are wired and how the environment influences the TRN structure.

### Abbreviations

*Hfx. volcanii*: (*Haloferax volcanii*)
TF: (transcription factor)
TRN: (transcriptional regulatory network)

## ACKNOWLEDGEMENTS

We thank the Duke University School of Medicine for the use of the Microbiome Core Facility, which provided NanoString service.

We thank the Duke University School of Medicine for the use of the DUGSIM, which provided ChIP-seq and RNA-seq service.

We thank Angie Vreugdenhil for her enthusiastic involvement in the creation of the Bioscreen analysis codeset.

## DATA AVAILABILITY

Whole genome sequencing data is accessible via NCBI Sequence Read Archive at PRJNA1003915. ChIP-seq and RNA-seq data are accessible at NCBI Gene Expression Omnibus via GSE240622. Code and raw input data used in analyses related to the study are provided in the GitHub repository at https://github.com/amyschmid/troR.

## CONTRIBUTIONS

Mar Martinez Pastor: Δ*arsR,* Δ*troR* (several intends with same results), *arsR*::HA strain; responsible for all the experimental designs, wet experiments and writing.

Saaz Sakrikar: RNA-seq and ChIP-seq analysis, preparation of supplementary tables.

Sungmin Hwang: Δ*idr,* Δ*sirR,* Δ*troR, idr*::HA, *sirR*::HA, *troR*::HA strains. Critical discussion during writing process. Manuscript editing.

Rylee Hackley: code for peak representation in figure 4 panel A.

Andrew Soborowski: whole genome sequencing analysis (breseq). Critical discussion about statistical parameters. Editing the manuscript.

Julie Maupin-Furlow: funding and guidance for construction of deletion strains; manuscript review and editing.

Amy Schmid: corresponding author.

## GUIDE TO SUPPLEMENTARY MATERIAL

**Figure S1.** Deletion of DtxR genes resulted in complete knockout strains.

**Figure S2.** Visualization of peaks detected across the *Hfx. volcanii* genome for each of TroR and SirR across replicate trials

**Figure S3.** Nanostring gene expression and B-galactosidase promoter-reporter experiments validating that deletion of *arsR* in the Δ*troR* background does not impact the expression of TroR-regulated gene targets.

**Table S1.** List of plasmids, strains, primers, and summary of whole genome sequencing analysis for deletion strains used in this study.

**Table S2.** List of peak annotations detected for SirR-HA binding by ChIP-seq

**Table S3.** TroR regulon data, including list of ChIP-seq peak annotations, RNA-seq data for genes differentially expressed in response to iron in the parent strain (H26), TroR cis-regulatory binding motif sequences detected throughout the genome, and annotations for genes in the TroR high confidence regulon.

**Table S4.** List of specific gene probes and resultant data from Nanostring gene expression analysis.

