## Supplemental Figure 1 for "TroR is the primary regulator of the iron homeostasis transcription network in the halophilic archaeon *Haloferax volcanii*"

### HVO_0819 (sirR)

| Predicted mutation | | | | | | |
|---|---|---|---|---|---|---|
| evidence | seq id | position | mutation | annotation | gene | |
| MC JC | NC_013967 | 739,215 | Δ692 bp | coding (4-695/696 nt) | *HVO_RS08630* → | metal-dependent transcriptional regulator |

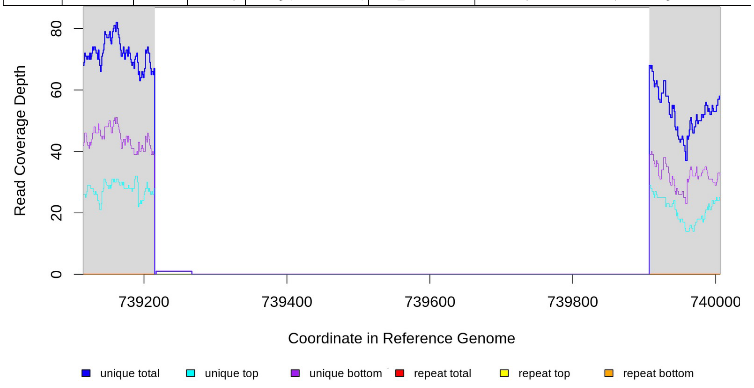


### HVO_0538 (idr)

| Predicted mutation | | | | | | |
|---|---|---|---|---|---|---|
| evidence | seq id | position | mutation | annotation | gene | |
| MC JC | NC_013967 | 468,633 | Δ431 bp | coding (2-432/435 nt) | *HVO_RS07290* ← | metal-dependent transcriptional regulator |

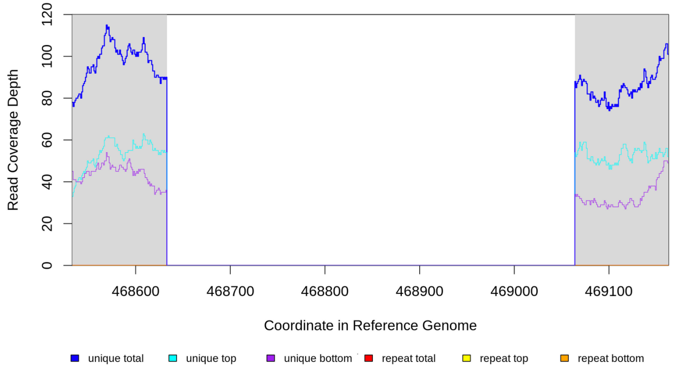


### HVO_0863 (troR)

| Predicted mutation | | | | | | |
|---|---|---|---|---|---|---|
| evidence | seq id | position | mutation | annotation | gene | |
| MC JC | NC_013967 | 778,851 | Δ453 bp | coding (1-453/453 nt) | *HVO_RS08835* → | metal-dependent transcriptional regulator |

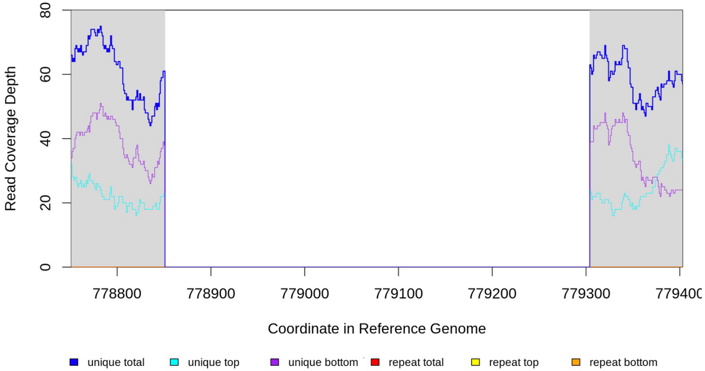


### HVO_1766 (arsR)

| Predicted mutation | | | | | | |
|---|---|---|---|---|---|---|
| evidence | seq id | position | mutation | annotation | gene | |
| MC JC | NC_013967 | 1,633,761 | Δ378 bp | coding (1-378/384 nt) | *HVO_RS13195* → | winged helix-turn-helix domain-containing protein |

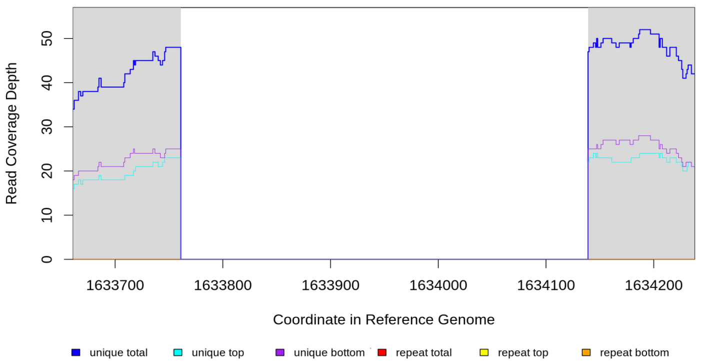


**Figure S1. Raw sequencing data for deleted gene loci.** In each panel, shown above is the results from breseq mutation analysis of whole genome sequencing data (see also details in Table S1). Below, a zoomed-in region of the whole genome resequencing data are shown for each of the three strains deleted for DtxR family paralogs. The Integrated Genome Viewer (IGV) genome browser was used to visualize the regions. Each zoomed region is labeled above with the name of the gene deleted. Colored lines are specified in the legends below each panel.
