## Supplemental Figure 2 for "TroR is the primary regulator of the iron homeostasis transcription network in the halophilic archaeon *Haloferax volcanii*"

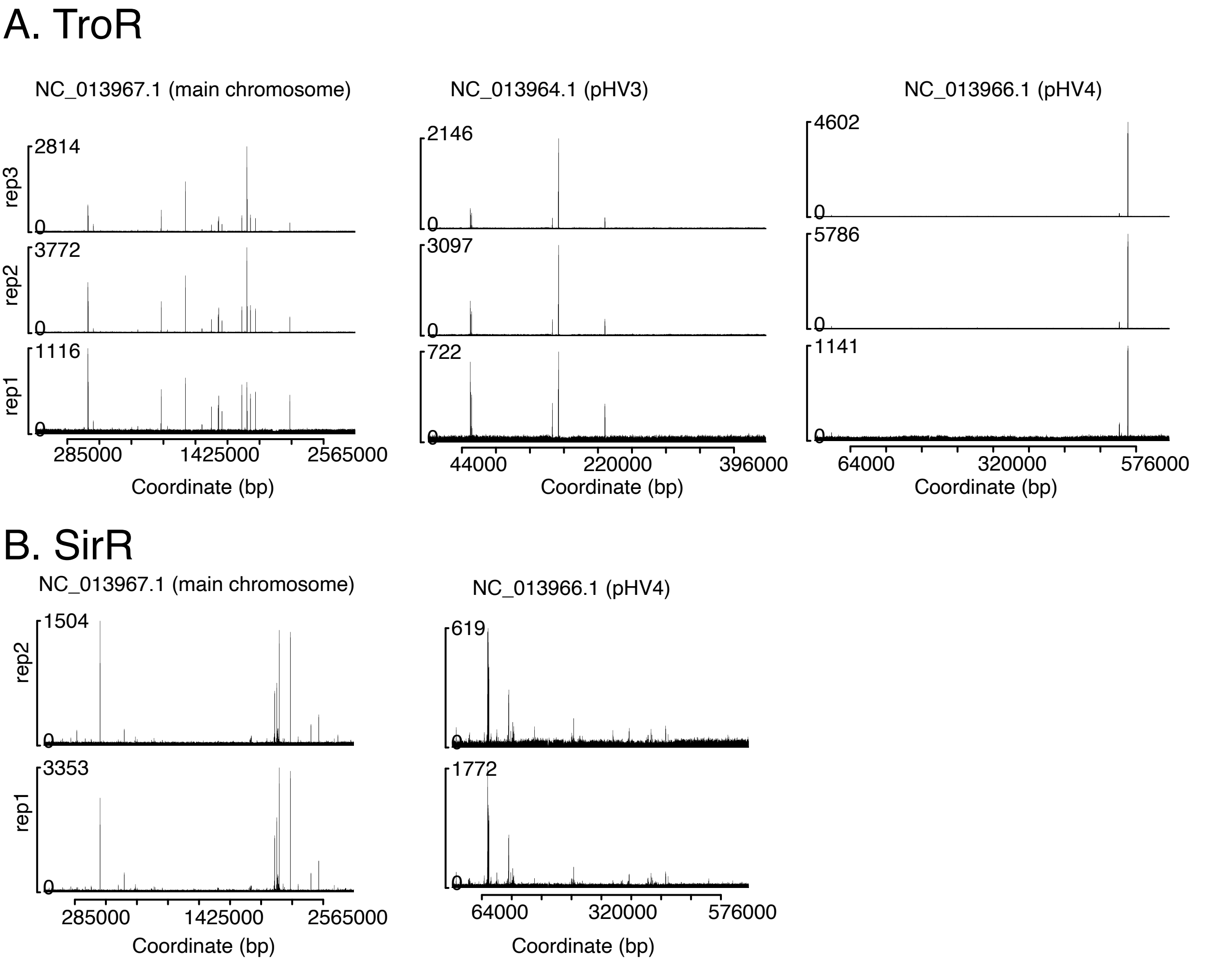


**Figure S2. All ChIP-seq peaks detected across replicates.** (A) Peaks detected for TroR-HA. Y-axis represents read depth, x-axis shows the genomic coordinate in base pairs (bp). Only chromosomal elements are shown where peaks were detected. Raw bam files of immunoprecipitation-enriched DNA bound to TroR were plotted for each of three replicates (rep1, rep2, rep3). The R package trackvieweR was used. (B) Peaks detected for each of two replicates for SirR-HA. Axes are as in (A). Raw bam files were plotted for IP samples using trackveiweR.
