## Supplemental Figure 3 for "TroR is the primary regulator of the iron homeostasis transcription network in the halophilic archaeon *Haloferax volcanii*"

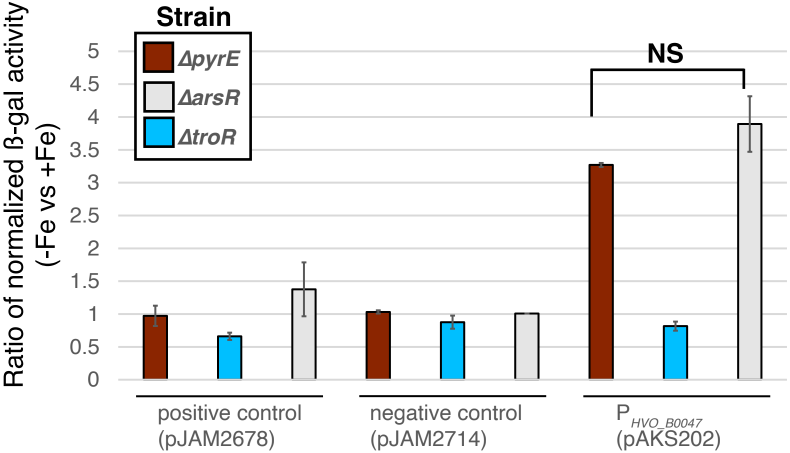


**Figure S3. ArsR does not co-regulate iron uptake genes with TroR.** ß-galactosidase promoter-reporter assay for the siderophore biosynthesis cluster promoter in the presence of iron (constructs pAKS2678, pAKS2714, and pAKS202, see also Table S1 and main text Figure 5). Ratio of β-gal activity in medium -Fe vs +Fe is compared between the Δ*pyrE*, Δ*troR*, and Δ*arsR* mutants. No significant difference was detected for the activity of this promoter in Δ*pyrE* vs Δ*arsR*. (note that $P_{HVO\_B0047}$ promoter activity ratio is constitutively de-repressed in both - and +Fe conditions, leading to ratio of ~1 in Δ*troR* (see also main text figure). Error bars represent standard error of the mean.
